# Widespread occurrence of the droplet state of proteins in the human proteome

**DOI:** 10.1101/2020.10.21.348532

**Authors:** Maarten Hardenberg, Attila Horvath, Viktor Ambrus, Monika Fuxreiter, Michele Vendruscolo

## Abstract

A wide range of proteins have been reported to condensate into a dense liquid phase, forming a reversible droplet state. Failure in the control of the droplet state can lead to the formation of the more stable amyloid state, which is often disease-related. These observations prompt the question of how many proteins can undergo liquid-liquid phase separation. Here, in order to address this problem, we discuss the biophysical principles underlying the droplet state of proteins by analyzing current evidence for droplet-driver and droplet-client proteins. Based on the concept that the droplet state is stabilized by the large conformational entropy associated with non-specific side-chain interactions, we develop the FuzDrop method to predict droplet-promoting regions and proteins, which can spontaneously phase separate. We use this approach to carry out a proteome-level study to rank proteins according to their propensity to form the droplet state, spontaneously or via partner interactions. Our results lead to the conclusion that the droplet state could be, at least transiently, accessible to most proteins under conditions found in the cellular environment.

**Significance:** Liquid-liquid phase separation of proteins results in biomolecular condensates, which contribute to the organisation of cellular matter into membraneless organelles. It is still unclear, however, whether these condensates represent a common state of proteins. Here, based on biophysical principles driving phase separation, we report a proteome-wide ranking of proteins according to their propensity to condensate into a droplet state. We analyze two mechanisms for droplet formation - driver proteins can spontaneously phase separate, while client proteins require additional components. We conclude that the droplet state, as the native and amyloid states, is a fundamental state of proteins, with most proteins expected to be capable of undergoing liquid-liquid phase separation via either of these two mechanisms.

---

It has been recently observed that proteins can self-assemble through a liquid-liquid phase separation process into a dense liquid phase, while maintaining at least in part their functional native states (1–4). These liquid-like assemblies of complex compositions are often referred to as biomolecular condensates, or membraneless organelles (1–4). Here, we refer to these dynamic and reversible condensates as droplets, in order to distinguish them from irreversible amyloids. Droplets can concentrate cellular components to perform efficiently a variety of different functions, with an increasing number of biological roles being discovered (1–4).

In this work, we investigate whether liquid-liquid phase separation can be expected to be a proteome-wide phenomenon. In this view, the condensation of proteins from the native state to the amyloid state may quite generally proceed through an intermediate dense liquid phase, which is typically metastable (5) (**Figure 1**). Different proteins may have different propensities to remain in this metastable phase, depending in particular on the free energy barrier between the droplet and amyloid states (**Figure 1**). This type of liquid-liquid phase separation is indeed typical of condensation phenomena (1, 6), and sometimes is referred to as the Ostwald step rule (7). One may think that for most proteins the free energy barrier between the droplet and fibrillar states is low, and therefore the droplet state cannot be readily observed (**Figure 1**). Indeed, this state may be difficult to detect due to a variety of reasons, including because experimental methods to probe its formation, in particular high-throughput ones, are still under development (8). Furthermore, our current understanding of the interactions that stabilize the metastable dense liquid phase is still incomplete.

**Figure 1.**
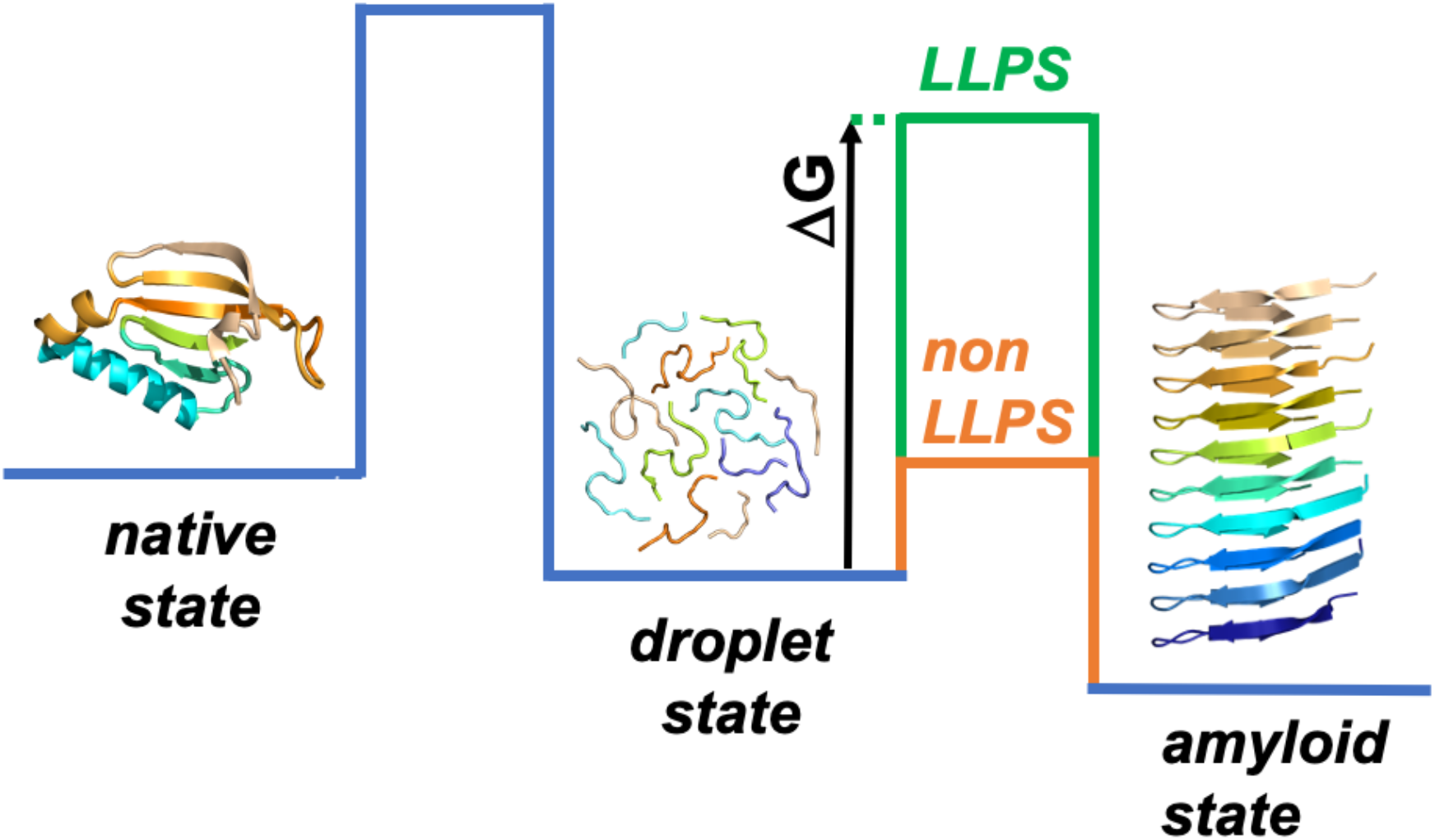
Liquid-liquid phase separation could be expected to be a proteome-wide phenomenon. Proteins that undergo condensation convert from the native state to the amyloid state through a dense liquid state (the droplet state). The stability of these different states (the minima in the free energy), as well as the conversion rates between them (the barriers in the free energy), are different for different proteins. For most proteins under cellular conditions, the native and droplet states could be expected to be metastable (58), being kinetically trapped by a free energy barrier (ΔG) between the droplet and fibrillar states. Proteins that can be observed in the droplet state tend to have a high free energy barrier (LLPS, green) while the other ones tend to have a low free energy barrier compared to the thermal energy (non-LLPS, orange). For certain proteins, the droplet state is functional, and it is stabilized by extrinsic factors, such as RNA and post-translational modifications.

Native and amyloid states are stabilized by specific interactions including hydrogen bonds, ionic interactions, and van der Waals contacts typical of ordered states and enthalpic in nature (9, 10). By contrast, in droplets, transient short-range aromatic cation-π and π-π, dipole-dipole, electrostatic and hydrophobic interactions have been observed, providing low-specificity, weak-affinity contacts characteristic of disordered states (11–16). These observations have lead to a series of prediction methods (11, 13, 17–19), which focused on specific side-chain interactions. The redundancy and multivalency of the interacting elements (20) suggests that conformational entropy is a driving force of the condensation (21), also including main chain contributions. Indeed, proteins exhibiting many binding configurations with a specific partner are often capable of forming droplets (22).

Here, we exploit the observation that many proteins exhibit high conformational entropy upon binding, which can be predicted from their amino acid sequences (23). Based on this result, we develop the FuzDrop method to predict the droplet-promoting propensity of proteins and their droplet-promoting profiles based on the conformational entropy of their free states and the binding entropy. Using this method, we identify a list of ‘droplet-driving’ proteins, which are predicted to undergo spontaneous liquid-liquid phase separation under physiological conditions, and estimate that they comprise about 40% of the human proteome. In addition, we also predict that about 80% of the proteins are ‘droplet-clients’, characterised by short droplet-promoting regions in their sequences, which facilitate condensation via interactions with suitable partners. Taken together, our results indicate that protein phase separation is a proteome-wide phenomenon.

## Results

### A framework to describe the interactions stabilising the droplet state

The premise of this work is that the droplet-state is characterized by low-specificity interactions and liquid-like conformational entropy. Thus, we hypothesized that proteins that are conformationally heterogeneous in their native states and maintain this property upon binding would be particularly prone to form the droplet state. In estimating the degree of conformational heterogeneity in both the native and bound states, we observe that proteins span a continuum between structural order and disorder (23, 24), which we will express by the probabilities of *p*_*D*_ (free state) and *p*_*DD*_ (bound state). We also note that interactions with high conformational entropy are realized via many different binding configurations, which can be achieved by both ordered and disordered domains (25). By contrast, ordered binding modes with low conformational entropy are mediated by well-defined interfaces, as exemplified by rigid docking or templated folding (26).

Ordered and disordered binding modes exhibit characteristic sequence signatures. Motifs mediating ordered binding modes have a strong compositional bias as compared to their embedding protein regions. In contrast, motifs mediating disordered binding modes are more similar to their flanking regions, which can be realized via a variety of sequence patterns and contact types, as their specificity stems from their distinct character as compared to their flanking regions (23). We have previously demonstrated (27) that by identifying such interaction elements based on compositional bias, it is possible to estimate structural order or disorder under cellular conditions in excellent agreement with *in vivo* proteomic studies (28).

### Properties of the proteins that can form the droplet state

#### Datasets of proteins representing the droplet state

We have analysed three public datasets of proteins reported to undergo liquid-liquid phase separation (see Methods). The first is the PhaSepDB dataset (http://db.phasep.pro) (29), which assembles data from three resources (Methods, **Table S1**): (1) proteins from the literature with *in vivo* and *in vitro* experimental data on liquid-liquid phase separation (REV, 351 proteins, Methods, **Table S1**), (2) proteins from UniProt associated with known organelles (UNI, 378 proteins, Methods, **Table S1**), and (3) proteins identified by high-throughput experiments of liquid-liquid phase separation (HTS, 2572 proteins, Methods, **Table S1**). The second dataset is PhaSePro (https://phasepro.elte.hu) (30), which identifies protein regions associated with liquid-liquid phase separation (PSP, 121 proteins, Methods, **Table S1**). The third dataset is LLPSDB (http://bio-comp.org.cn/llpsdb) (31), which assembles proteins observed to undergo *in vitro* liquid-liquid phase separation with well-defined experimental conditions and phase diagrams (Methods, **Table S1**). LLPSDB distinguishes whether proteins can phase separate spontaneously as one component (droplet-driving proteins, LPS-D, 133 proteins, Methods, **Table S1**) or require a partner to undergo liquid-liquid phase separation (droplet-client proteins, LPS-C, 41 proteins, Methods, **Table S1**). In this dataset, 77 proteins exhibit both droplet-driving and droplet-client behaviors.

To create a dataset for liquid-liquid phase separation, we merged the proteins in the REV, PSP and LPS-D datasets, which we consider as drivers of droplet formation (453 unique proteins, LLPS dataset, Methods, **Table S1**). We generated two negative control datasets, one with human proteins only and another with a mixture of organisms (**Table S2**). For the human negative set (hsnLLPS dataset, 18108 proteins, Methods) we excluded from the Swiss-Prot human proteome all proteins that appeared in any of the liquid-liquid phase separation datasets (REV, UNI, HTS, PSP, LPS-D, LPS-C) (29–31) (**Table S2**). For the negative set corresponding to multiple organisms (nsLLPS, Methods), we derived the organism distribution from the LLPS dataset. To build a control dataset, we considered organisms populated more than 1% in the LLPS dataset and used their proteomes from UniProt (*Caenorhabditis elegans, Chlamydomonas reinhardtii, Drosophila melanogaster, Homo sapiens, Mus musculus, Rattus norvegicus, Saccharomyces cerevisiae, Schizosaccharomyces pombe, Xenopus laevis*, Methods) and removed all proteins present in the LLPS or HTS datasets. We then randomly chose sequences according to the frequencies of these organisms in the LLPS dataset with 10 times enrichment (nsLLPS, **Table S2**, Methods).

#### Analysis of the amino acid compositions of droplet-driving and droplet-client proteins

Droplet-driving proteins are enriched in disorder-promoting residues (P, G, S) and depleted in order-promoting (F, I, V, C, W) residues as compared to non-phase separating proteins (**Figure 2A**). N and Q, which are distinguished in prion-like domains (32) are more abundant in droplet-driving proteins than those, which were not reported to undergo LLPS. However, droplet-driving proteins are not significantly enriched in residues that mediate π-π and cation-πinteractions (Y, R), as compared to non-phase separating (nsLLPS) proteins (**Figure 2A, Table S3**). These results indicate that droplet-formation does not depend on a specific contact type, but it can rather be realized in many ways by low-specificity interactions. The composition of droplet-driving proteins is in between that of globular proteins (33) and disordered proteins in the DisProt database (33), they are more abundant in order-promoting residues (W,C,Y,F, I,V) as compared to disordered proteins (**Figure S1B**) and enriched in disorder-promoting residues (P,D,E) as compared to globular proteins (34) (**Figure S1A**). Aromatic residues observed in disordered regions, for example in nucleoporins, often mediate low-affinity interactions (35). These compositional properties reflect the preference of droplet-driving proteins for the disordered state in the bound form, which is comparable to protein complexes with disordered binding modes (23).

**Figure 2.**
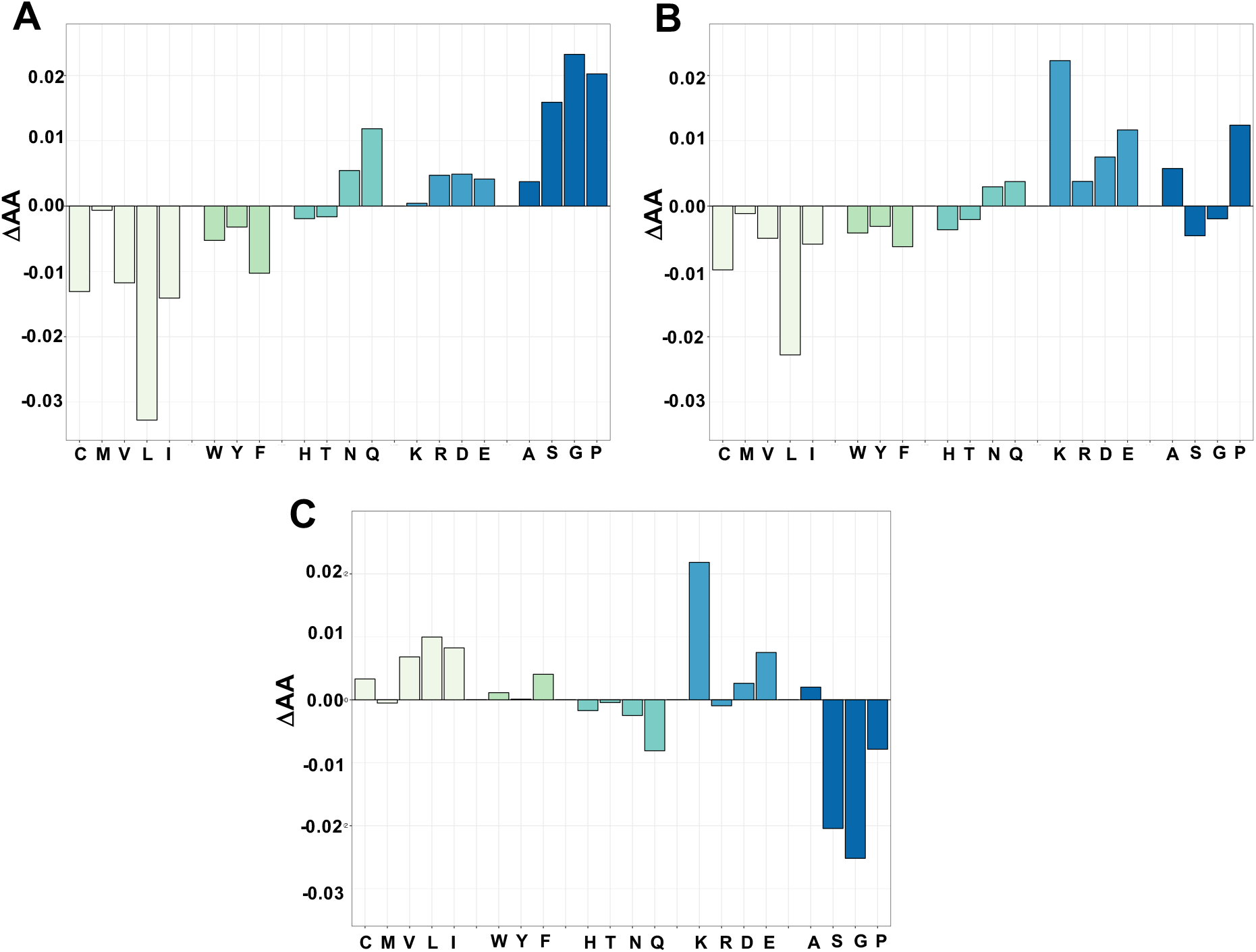
Differential amino acid compositions of droplet-driver and droplet-client proteins. **(A)** Differences in amino acid compositions (ΔAA) of droplet-driver proteins in the LLPS dataset and of proteins not reported to phase separate (nsLLPS). **(B)** Differences in amino acid compositions of droplet-client proteins that require additional components for phase separation (LPS-C dataset) and of proteins that have not been reported to phase separate (nsLLPS). **(C)** Differences in amino acid compositions of droplet-client proteins (LPS-C) and droplet-driver proteins (LLPS). Amino acids grouped as hydrophobic (light green), aromatic (green), hydrophilic (turquoise), charged (steel blue), disorder-promoting (dark blue) (34). The standard errors and the significances of the differences by Kolgomorov-Smirnov test are shown in **Table S3**.

As compared to non-phase separating proteins, droplet-client proteins are enriched in charged residues (D,K,E) and disorder-promoting prolines (**Figure 2B, Table S3**). Droplet-client proteins exhibit characteristic differences from droplet-driving proteins, as they are enriched in charged residues (K, E), and hydrophobic motifs (L, V, I), while being depleted in amyloid-promoting (N, Q), phosphorylation-promoting (S) and disorder-promoting (G) residues (**Figure 2C, Table S3**). The amino acid composition of droplet-clients is thus more similar to structured than disordered proteins (**Figure S1B**).

#### Analysis of the conformational entropy of droplet-driving and droplet-client proteins

We observed that different protein datasets representing the droplet state have markedly different characteristics in their conformational entropy in the free state and its change upon binding. Drivers of droplet formation (LPS-D) have high level of disorder in free (*p*_*D*_) and bound states (*p*_*DD*_), while droplet clients (LPS-C) are mostly ordered in both forms (**Figure 3A,B**). Proteins in the REV and PSP datasets exhibit disordered binding modes, which are comparable to droplet-driver proteins, so they likely phase separate spontaneously. Proteins associated with known membraneless organelles (UNI), or identified by high-throughput experiments (HTS) (29) have significantly lower conformational entropy in both free and bound states, thus likely have components, which form droplets via partner-interactions. Comparison of spontaneously phase-separating and non-phase separating proteins (**Figures 3C,D**) indicates that a high conformational entropy is a characteristic of the droplet state.

**Figure 3.**
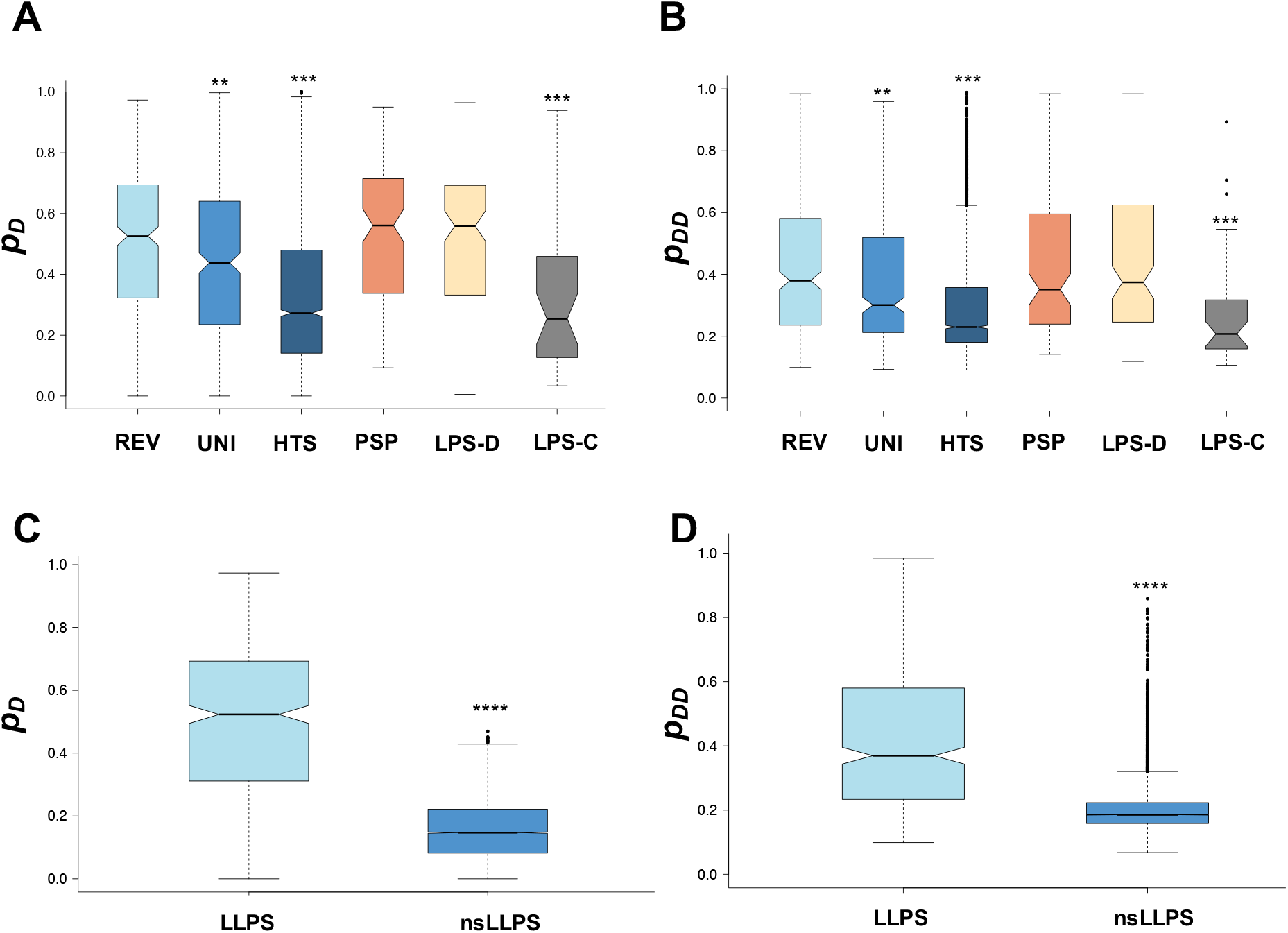
Conformational properties in different datasets of LLPS proteins in the free and bound states. PhaSepDB literature reviewed (REV, light blue), PhaSepDB human organelle associated proteins from UniProt (UNI, steel blue), PhaSepDB proteins identified by high-throughput experiments (HTS, dark blue), PhaSePro (PSP, orange), LLPSDB one-component proteins (droplet-drivers, LPS-D, wheat) and two-component (droplet-clients, LPS-C, gray) phase separating proteins. **(A)** The probability for the disordered state (p_D_) in the free form was characterized by the fraction of disordered residues, as computed by the Espritz NMR program (36). Residues are classified as disordered if they have an ID score ≥ 0.3089. The fraction of disordered residues was computed per protein as (N_ID_/N_AA_) and these values were averaged for each dataset. **(B)** Probability for disordered binding (p_DD_) was computed by the FuzPred program (23). The median p_DD_ value was determined for each protein and these values were averaged for each dataset. **(C-D)** Comparison of *p*_*D*_ and p_DD_ in droplet-driving (LLPS, light blue) and non-phase separating proteins (nsLLPS, dark blue). Statistical significances were computed by Mann-Whitney test using the R program (** p < 10^−3^, *** p< 10^−5^, **** p< 10^−10^).

### Sequence-based prediction of droplet propensity profiles of proteins

Based on the analysis reported above, in this section we present a method of predicting the sequence-based profile of the propensity of proteins to form spontaneously the droplet state (*p*_*DP*_). To achieve this result, we define the probability of residue *A*_*i*_ to be involved in spontaneous phase separation by *p*_*DP*_ (*A*_*i*_) using a binary logistic model as

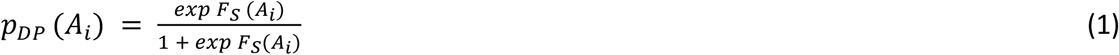

where *F*_*S*_ (*A*_*i*_) is the scoring function for the residue

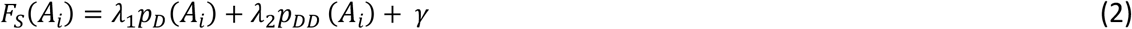

where *p*_*D*_(*A*_*i*_) is the probability for the disorder in the free state, and *p*_*DD*_(*A*_*i*_) is the probability for disordered binding (23). *p*_*D*_(*A*_*i*_) contains an estimate of the conformational entropy in the unbound form, while *p*_*DD*_(*A*_*i*_) contains an estimate of the binding entropy. *λ*_1_ and *λ*_2_ are the linear coefficients of the predictor variables and *γ* is a scalar constant (intercept), which were determined using the binary logistic model (Methods, **Table S4**). *p*_*D*_ was derived from the disorder score as computed using the Espritz NMR algorithm (36), with the best performance on disordered protein complexes (23). The *p*_*DD*_ values were predicted by the FuzPred method, which describes binding modes under cellular conditions (27). The *p*_*D*_ and *p*_*DD*_ values capture the balance between enthalpy and entropy that stabilises the droplet state, which is associated with the non-specific nature of a variety of side-chain interactions.

To train our model, we used a dataset of droplet-promoting regions, with an evidence to mediate spontaneously phase separation (Methods, **Table S1**). As a negative set, we defined regions in non-phase separating proteins with the same length distribution as in the positive set (**Methods**). The size of the negative set was 10 times that of the positive set and we applied stratified sampling in the training. We found that the linear coefficients were robust over many random selections of the positive and negative sets, as well as the training set size (**Table S4**). In the final parametrisation, the linear coefficients of both disorder and binding mode were positive, reflecting the preference for a disordered bound state in the droplets. The threshold to mediate droplet-formation was derived from the binary logistic model (*p*_*DP*_ ≥ 0.60).

To estimate the performance of the method we calculated an AUC value of 87.0% on the training set (0.8 of the droplet dataset) and an AUC value of 85.9% on the test set (**Table S4**, Methods). We applied these coefficients on all droplet regions and obtained an AUC value of 84.4%. These results illustrate that the parameters are robust across droplet regions from different organisms. We also note that droplet-promoting propensity profiles of proteins that were observed to form droplets under cellular conditions and those that were detected only by in vitro experiments are not significantly different (**Figure S3**).

We have thus developed the FuzDrop method to predict droplet-promoting propensities of residues from the primary sequence based on the conformational entropy of disordered binding modes in droplets.

### Droplet-promoting propensity profiles of TDP-43 and α-synuclein

We applied the FuzDrop method to predict the droplet-promoting propensities of two proteins reported to undergo liquid-liquid phase separation, TDP-43 (37) and α-synuclein (38, 39). Our results indicate that the low-complexity (LC) region of TDP-43 (residues 262-414) mediates spontaneous phase separation. We note that the α-helical segment (residues 320-331), which constitutes the amyloid core in TDP-43 fibrils (40) (**Figure 4A**) and is predicted to undergo disorder-to-order transition upon binding, also has a high droplet-promoting propensity (**Figure 4A**).

**Figure 4.**
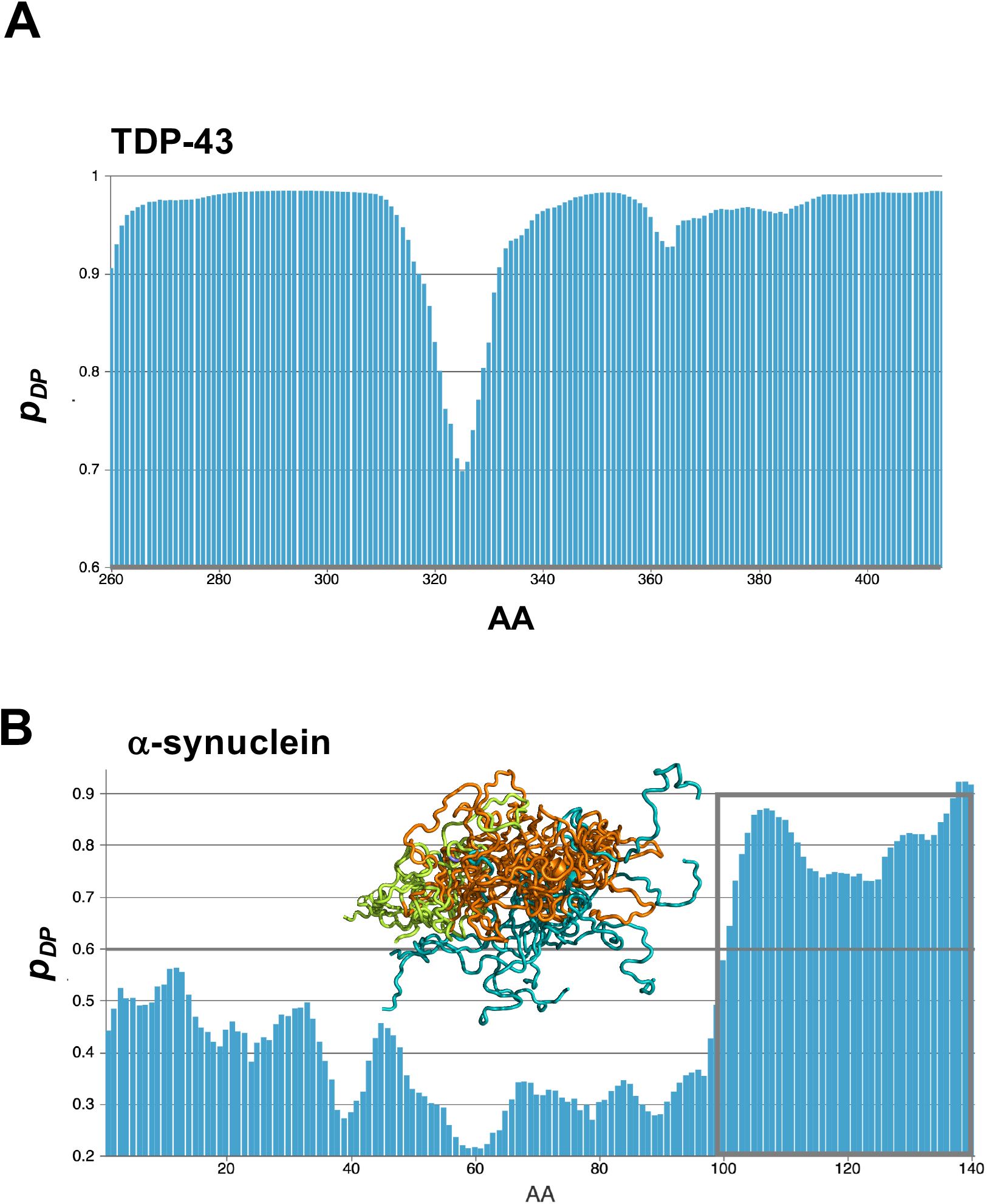
Droplet-promoting propensity profiles (*p*_*DP*_) of the TDP-43 LC domain and of α-synuclein. (**A**) The TDP-43 low complexity (LC) domain has overall high droplet-promoting propensities. The depletion in the droplet profile corresponds to the α-helical segment (orange), which is involved in the amyloid core. The N-(lime) and C-(blue) flanking regions are disordered in the NMR structure of the G335D mutant (PDB:2n4g). **(B)** The disordered C-terminal region of α-synuclein (blue) is predicted to drive droplet formation. The N-terminal region (lime), which folds into α-helix has intermediate *p*_*DP*_ values. The ensemble is derived from Protein Ensemble Database (PED9AAC). The p_LLPS_ threshold is indicated by bold grey line.

In the case of α-synuclein, the highly disordered C-terminal region (residues 98-140), which also remains disordered upon binding to lipid vesicles (41), is predicted to drive the formation of the droplet state, (**Figure 4B**). The central NAC region has lower *p*_*DP*_ propensity to spontaneously phase separate, but may be involved in droplets via hydrophobic protein interactions, which are absent from β-synuclein and γ-synuclein (38).

### Sequence-based prediction of droplet-driving proteins

In this section, we present a method of ranking proteins according to their propensity to form the droplet state. In order to achieve this result, we estimate the probability of liquid-liquid phase separation (*p*_*LLPS*_) using a binary logistic model (Methods) with a scoring function (*F*_*LLPS*_) derived from residue droplet-promoting propensities and a term for hydrophobic interactions

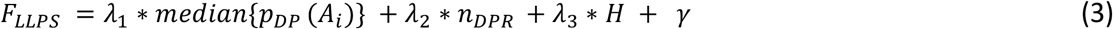

where *median*{*p*_*DP*_(*A*_*i*_) median of the residue droplet-promoting propensities, *n*_*DPR*_ is the number of long droplet-promoting regions (DPRs, ≥25 consecutive residues with p_DP_ ≥ 0.6) and H is a hydrophobic term (≥6 residue hydrophobic motifs within disordered regions, Methods). *λ*_1_, *λ*_2_ and *λ*_3_ are the linear coefficients of the predictor variables and *γ* is a scalar constant (intercept), which we determined on the LLPS_train_ and nsLLPS_train_ datasets (Methods, **Table S5**). We found that the linear coefficients were robust over many random selections of the positive and negative sets, as well as the training set size (**Table S5**). The threshold to mediate spontaneous liquid-liquid phase separation was derived from the binary logistic model (*p*_*LLPS*_ ≥ 0.61). We propose that the *p*_*LLPS*_ values express the droplet-driving potential under physiological conditions, as droplet-promoting propensities of proteins that form droplets under physiological conditions and those that were detected to phase separate only in vitro do not deviate significantly (**Figure S2**). We also note that using non-physiological conditions, such as high concentrations of protein and crowding agents, can induce liquid-liquid phase separation at *p*_*LLPS*_ values below the threshold, especially if droplet-promoting regions are present.

To estimate the performance of the method we calculated an AUC value of 88.3% on the training set (0.75 of the LLPS dataset) and an AUC value of 90.7% on the test set, using stratified sampling (**Table S5**, Methods). As an attempt to further improve performance, we incorporated a π-πterm (19) into the scoring function of the logistic model (Methods). Adding this term, slightly increased the performance of the model (AUC=92.2%, **Table S5**) with a moderate contribution to the scoring function. These results are in accord with the presence of π-πinteractions in many droplet proteins, but also show that these interactions are not pre-requisites for droplet formation.

The performance and robustness of the model (Eq. 3, **Table S5**) demonstrates that the droplet state can be predicted from sequence based on the estimated conformational entropy of binding and a non-specific enthalpy term. We also note, that our model by Eq. 3 serves as a general framework for predicting droplet-driver proteins. Accumulating data collected using more systematic and uniform experimental approaches (8) will enable further refinement of the parameters in our model and to predict the minimum concentration for phase separation, although this property can be expected to be highly dependent on the cellular conditions.

### Region specificity of the FuzDrop method and experimental validation of the predictions

We note that estimates of the overall propensity of a protein to form the droplet state cannot be readily obtained by a simple average of the values of the profiles of Eq. 2. This overall propensity is also determined by specific regions, rather than only by the general properties of the entire sequence, including in particular droplet-promoting regions and short motifs within disordered regions, which are prone to establish hydrophobic interactions (Eq. 3). This point can be illustrated by distinct behaviours of α-synuclein and β-synuclein (38). The C-terminal region of both proteins possess a droplet-promoting region, with a preference for disordered binding modes (**Figures 4 and S3**). In addition, the NAC region of α-synuclein contains 8 hydrophobic residues, biased for disordered binding, which can exert a non-specific driving force (resembling hydrophobic collapse) for droplet formation. Notably however, β-synuclein and γ-synuclein, which lack these residues (**Figure S3**), were not observed to undergo liquid-liquid phase separation under physiological conditions (38).

The predicted *p*_*LLPS*_ values by the FuzDrop method (0.62 for α-synuclein, 0.54 for β-synuclein and 0.40 for γ-synuclein) suggest that β-synuclein and γ-synuclein have lower propensity to adopt the droplet state as compared to α-synuclein. Indeed, γ-synuclein did not phase separate under any of the experimental conditions tested (38). To validate the predictions close to the prediction threshold, we explored β-synuclein phase behavior in a set of *in vitro* experiments (**Figure 5**). In line with previous observations (38, 39), we did not observe any droplets after incubating high concentrations of FITC-labelled β-synuclein on a glass surface, whereas we did observe droplets for FITC-labelled α-synuclein (**Figure 5A,B**). As hydrophobic effects are important for α-synuclein droplet formation, and considering that β-synuclein lacks the predominantly hydrophobic segment in the NAC region, we reasoned that raising the experimental temperature would increase the strength of residual hydrophobic interactions, allowing the protein to cross the phase barrier. Indeed, β-synuclein formed micrometer sized droplets when incubated at 30 °C and at high concentrations (**Figure 5C**). Droplets formed by FITC-β-synuclein were initially liquid-like, as evidenced by fast recovery after photobleaching (FRAP), but showed rapid conversion to a gel-like state (**Figure 5D**). The phase-separation behaviour of β-synuclein illustrates that protein phase separation is highly dependent on the experimental conditions, that proteins with a predicted *p*_*LLPS*_ below the threshold (*p*_*LLPS*_ ≥ 0.61) require more extreme conditions to adopt the droplet state, and that the droplet state of these proteins is generally short-lived.

**Figure 5.**
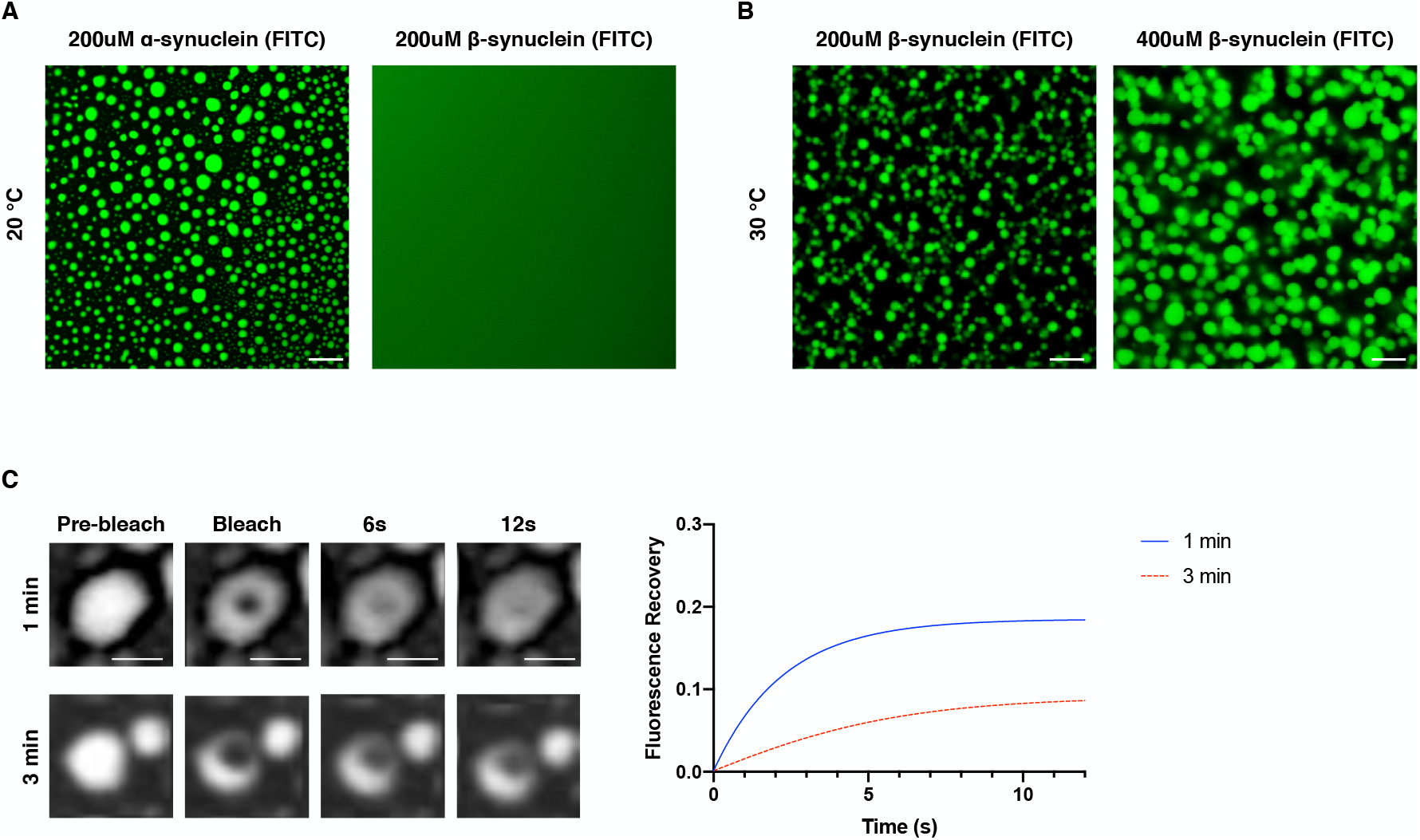
Region specific phase behaviour of α-synuclein and β-synuclein. (**A**) FITC-labelled β-synuclein (*p*_*LLPS*_ 0.54), which lacks the characteristic NAC region found in α-synuclein, does not phase separate at high concentrations (200 μM) and under crowding conditions (10% w/v PEG), whereas FITC-labelled α-synuclein (*p*_*LLPS*_ 0.62) readily forms droplets under the same conditions. (**B**) Increasing the experimental temperature to 30 °C does lead to rapid coalescence of β-synuclein into micrometer sized droplets. (**C**) Rapid fluorescence recovery after photobleaching (FRAP) of a small area within a droplet (*top*) 1 minute after phase separation; FRAP 3 minutes after phase separation (*bottom*); non-linear fit of fractional fluorescent recovery over time (*right*). Scale bar: 10 μm (**A**) and 5 μm (**C**).

As an additional test of our predictions, we ranked a set of proteins associated with Alzheimer’s disease (42) based on their predicted FuzDrop scores (**Table S6**) and selected one of the top candidates, complexin-1 to experimentally test our predictions (**Figure 6**). To assess whether complexin-1 can form droplets through liquid-liquid phase separation, we incubated Alexa-488 labelled complexin-1 on a glass surface under crowding conditions at physiological pH (**Methods**). After a brief lag phase (< 1 minute), complexin-1 formed micrometer-sized droplets in suspension (**Figure 6A**). The droplets where characteristic of a liquid phase, as they showed distinct wetting behavior after prolonged incubation (>10 minutes) (**Figure 6A**) and fused upon making contact (**Figure 6B**). Furthermore, molecules within the droplets showed local rearrangement, as evidenced by rapid fluorescent recovery after photobleaching (FRAP) (**Figure 6C**). We also predicted that disordered N-terminal region of complexin-1 drives its liquid-liquid phase separation (**Figure S4**). This region cooperatively interacts with the SNARE complex and plasma membrane (43) to facilitate synaptic vesicle fusion (44). Phase separation may contribute to activation of complexin-1 by relieving its autoinhibition, which is a common mechanism by the droplet state (21).

**Figure 6.**
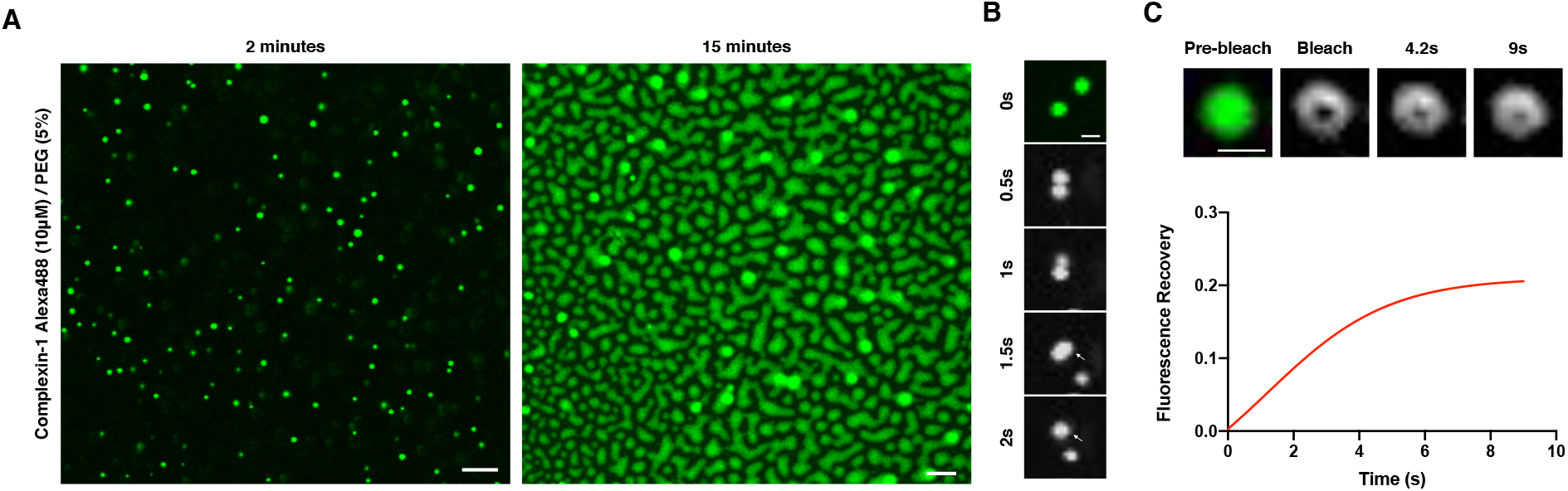
Complexin-1 undergoes liquid-liquid phase separation. (**A**) Alexa488-labelled complexin-1 (10 μM) coalesces into micrometer-sized droplets under crowding conditions (*left*). Droplets exhibit a wetting phenotype when encountering a glass surface (*right*). (**B**) Complexin-1 droplets readily fuse when in close proximity (<1 μm) and relax into a round structure after fusion. (**C**) Rapid fluorescence recovery after photobleaching (FRAP) of a small area within a droplet (*top*); non-linear fit of fractional fluorescent recovery over time (*bottom*). Scale bar: 5 μm (**A**) and 1 μm (**B, C**).

### Droplet-driving and droplet-client proteins in the human proteome

We applied the prediction method to estimate the proteins capable of undergoing spontaneous liquid-liquid phase separation (droplet-driving proteins) in the Swiss-Prot human proteome. We thus ranked the proteins in the human proteome according to their propensity to form the droplet state (**Table S7**), and estimated that about 40% of them are capable of spontaneous droplet formation.

This list contains only about 60% of the human proteins currently associated with membraneless organelles (UNI). This fraction is even lower for proteins identified by high-throughput experiments (HTS), including organelle purification (45, 46), affinity purification (47, 48), immunofluorescence image-based screen (49, 50), and proximity labelling (51, 52) (**Figure S5**). As the FuzDrop approach was developed for proteins that drive droplet formation, our results indicate that membraneless organelles contain also proteins that undergo phase separation by being driven by a partner (droplet-client proteins). We observed that droplet-clients have a lower conformational disorder in both free and bound states (**Figure 3**), suggesting the involvement of distinguished, local motifs. Thus, the droplet-client mechanism can provide a route for structured proteins to be engaged in condensates via specific droplet-promoting regions.

To investigate the properties underlying the droplet-client mechanism, we analyzed the presence of long and short droplet-promoting regions in the droplet-driver (LLPS) and droplet-client (LPS-C) datasets (**Table 1**). We found that ~90% of droplet-client proteins contain a short droplet-promoting region (≥10 residues), while only ~60 % have a long ones (≥25 residues). The frequency of short and long droplet-promoting regions in proteins, identified by high-throughput experiments is comparable to droplet-client proteins (**Table 1**), indicating that they follow a partner-induced client mechanism. In contrast, the frequency of droplet-promoting regions in proteins associated with human membraneless organelles is comparable to droplet-drivers (**Table 1**). Considering their lower droplet-promoting propensities (**Table S7**), these results indicate that proteins in membraneless organelles likely follow both driver and client mechanisms.

**Table 1.** Percentage in different datasets of proteins containing regions predicted to be droplet-promoting. Droplet promoting regions were identified with the number of consecutive residues with p_DP_ ≥ 0.6, either ≥ 25 (column 2) or ≥ 10 (column 3). The list presented are: membraneless organelles (UNI), high-throughput experiments (HTS), droplet drivers (**Table S1**, LLPS) and droplet clients (**Table S1**, LPS-C).

| | $p_{\text{DPR}} \geq 25$ residues<br>(%) | $p_{\text{DPR}} \geq 10$ residues<br>(%) |
| --- | --- | --- |
| Homo sapiens | 62 | 83 |
| Membraneless organelles | 84 | 94 |
| High-throughput experiments | 65 | 87 |
| Droplet-drivers | 87 | 94 |
| Droplet-clients | 60 | 88 |

Overall, we thus estimate that over 80% of the proteins in the human proteome contain regions that can mediate droplet formation. Half of these proteins can condensate spontaneously, while the other half can do so by interacting with other components (**Table 1**). We have also observed that the number of droplet-promoting regions is comparable in proteins, which were observed to form droplets under physiological conditions, and those, which detected by *in vitro* experiments (**Figure S2**), corroborating the relevance of the predictions under cellular conditions. We then extended these results to other organisms (**Table S8**), leading to the suggestion that the droplet-state is a proteome-wide phenomenon.

## Discussion and Conclusions

Increasing evidence indicates that a wide range of proteins unrelated in sequence, native structure and function can form biomolecular condensates (1, 2, 4, 53). These observations suggest that the droplet state may have a generic nature and be accessible to most proteins.

This possibility may not be immediately evident from the data currently available because the condensation of different proteins has been reported for experimental conditions often far from the physiological ones. Moreover, a full understanding of the interactions driving droplet-formation has not been achieved yet, owing to a wide variety of sequence motifs, which have been associated with the droplet state.

In this work, we have exploited that a large fraction of the proteins in the human proteome have positive binding entropies by visiting an ensemble of bound states (54, 55), which is realized via disordered binding modes. We thus hypothesized that the high conformational entropy associated with non-specific side-chain interactions contributes to the stabilization of the droplet state, and generated a model to quantify it from sequence. We have shown that droplet-promoting propensities can be predicted using such generic model without the explicit incorporation of specific types of interactions. The specificity of our model originates in local, compositional sequence biases, which are used to estimate the entropy in the bound state (23). That is, both hydrophobic or hydrophilic motifs can selectively mediate interactions if they are embedded in an environment of opposite character, explaining how selectivity can be achieved via a wide variety of interactions and contact types. We have shown earlier that this approach is capable to describe ordered and disordered binding under cellular conditions (27).

Using these general principles, we developed the FuzDrop method to predict droplet-promoting profiles and propensity of proteins to drive droplet formation. Applying this prediction method to different datasets of phase separating proteins, we described two mechanisms of droplet-formation: (1) The droplet-driver mechanism, which does not require addition components to phase separate mostly relies on high conformational entropy of the full protein sequence, and (2) The client mechanism, which is induced by protein interactions appears to more local, and dependent on a droplet-promoting region. Our results indicate that proteins may use the driver or the client mechanisms, or a combination of them, to form droplets.

Our proteome-wide analysis indicates that the presence of droplet-promoting regions is widespread in the sequences in the human proteome. Based on this analysis we conclude that the droplet state is accessible, even if only transiently, for most proteins. In ~40% of human proteome it is predicted to occur spontaneously, whereas an approximately equal fraction may require a variety of cellular components or non-physiological conditions. Proteins in known membraneless organelles represent a combination of these mechanisms, whereas those identified by high-throughput studies mostly represent droplet-clients.

Taken together, these results indicate that the droplet state is likely to be a fundamental state of proteins, alongside the native and amyloid states.

## Materials and Methods

### Datasets of phase separating proteins

All data in the present study were downloaded from public datasets without modifications (**Table S1**). The REV, UNI and HTS datasets were assembled from the PhaSepDB dataset (http://db.phasep.pro) (29). The 351 proteins in the REV dataset were collected based on curated literature search; the 378 proteins in the UNI dataset were associated with human organelles in UniProt; the 2572 proteins in the HTS dataset were identified in high-throughput experiments. The PSP dataset contained 121 proteins from PhaSePro database (https://phasepro.elte.hu) (30) with regions involved in LLPS identified. 174 proteins in the LLPSDB dataset (http://bio-comp.org.cn/llpsdb) (31) dataset were observed to undergo *in vitro* liquid-liquid phase separation for which the experimental conditions were also specified. All proteins, which were observed to form droplet spontaneously were assigned to the LPS-D dataset and only those, phase separation of which was dependent on interactions with a partner were in LPS-C dataset. The LLPS dataset contained 453 non-redundant proteins, by merging the REV, PSP and LPS-D datasets (**Table S1**). 144 regions, which were identified to mediate droplet formation were assembled from the PhaSePro dataset (30), and were grouped based on the evidence for spontaneous, or partner-assisted phase separation in the LLPSDB dataset (31) (DPR, **Table S1**).

### Datasets of non-droplet forming proteins

All human proteins included in the phase separation datasets (LLPS, and LPS-C) were removed from the Swiss-Prot human proteome, resulting in 18,108 sequences (hsnLLPS, **Table S2**). We also generated negative set for phase separation (nsLLPS, **Table S2**), which reflected the composition of the LLPS dataset using organisms represented >1% in the LLPS dataset (*Caenorhabditis elegans, Chlamydomonas reinhardtii, Drosophila melanogaster, Homo sapiens, Mus musculus, Rattus norvegicus, Saccharomyces cerevisiae, Schizosaccharomyces pombe, Xenopus laevis)*. Only Swiss-Prot sequences were used except for *Xenopus laevis*. Sequences were randomly chosen from these pools to match their frequency in LLPS. The size of the nsLLPS dataset was 10 times more than that of the LLPS dataset.

#### Analysis of amino acid compositions

The properties of LLPS proteins were compared to proteins with disordered regions in the Disprot v7 database (34) and the composition of globular proteins from the Protein Data Bank (33). We used a bootstrap approach to compare the amino acid compositions in proteins of the LLPS, LPS-C (**Table S1**) and nsLLPS (**Table S2**) datasets and the statistical significance of the pairwise differences were determined by a two-sample Kolmogorov-Smirnov test of the R program (**Table S3**). We also computed the absolute maximum distances between the cumulative distribution functions (**Table S3**). Standard error (SE) was calculated as *SE* = *SD*/√*n*, where *SD* represents the standard deviation of the bootstrapped differences and n represents sample size.

#### Predicting residue droplet-promoting propensity

##### Binary logistic regression model

Droplet-promoting propensity (*p*_*DP*_) was defined as a probability of a binary response, whether a residue can promote spontaneous phase separation or not. We used two predictor variables (Eqs. 1 and 2): i) probability of disorder in the free state (*p*_*D*_) was predicted by the Espritz NMR program (36). ii) the probability of disordered binding (*p*_*DD*_) and was computed by the FuzPred program (23). These two quantities approximated the conformational entropy in free state and its change upon binding.

### Training and parametrisation

As a positive set, we used 67 droplet-promoting regions, with an evidence for mediating spontaneous phase separation (**Table S1**). As a negative set, we randomly chose regions from proteins in 9 representative organisms (*Caenorhabditis elegans, Chlamydomonas reinhardtii, Drosophila melanogaster, Homo sapiens, Mus musculus, Rattus norvegicus, Saccharomyces cerevisiae, Schizosaccharomyces pombe, Xenopus laevis)* without an evidence to spontaneously form droplets or serve as droplet clients. Frequencies of proteins were set according to the droplet dataset (**Table S1**); with a length distribution matching that of the positive droplet-promoting region (DPR) dataset. The size of the negative set was 10 times of the positive set and we applied stratified sampling.

We used the R program to determine the coefficients the independent variables (*p*_*D*_ and *p*_*DD*_, Eqs. 1 and 2) on the training set, which was chosen as 0.6-0.8 of the positive DPR set. The performance of the different models was evaluated based on area-under-the-curve (AUC), specificity, sensitivity and accuracy, which were computed by the R program (**Table S4**). Owing to the length-dependence of the characteristics of the droplet-promoting regions, we used the coefficients, which were obtained for regions < 200 residues. The threshold for droplet-promoting propensity (*p*_*DP*_ => 0.5994) was determined based on the logistic model, and were in good agreement for the training and test sets.

#### Predicting the propensity of proteins to drive droplet-formation

A binary logistic model was used to estimate the probability of a binary response, whether protein spontaneously forms droplets or not, based on three predictor variables (Eq. 3): The median of the residue-based p_DP_ values, the number of droplet-promoting regions (n_DPR_ => 25 residues) and a factor representing weak hydrophobic interactions. To distinguish between hydrophobic interactions driving structure formation and those in droplets, we used hydrophobic motifs (=> 6 consecutive residues), which were located in disordered regions. The threshold was set to −1.3 based on the Kyte-Doolittle hydrophobicity scale, to include S, T and Y capable to undergo phosphorylation. As data regarding droplet-forming proteins is rapidly expanding we aimed to use a model, which is general and can be re-optimised if more and more specific information will be available.

### Training and parametrisation

We divided the positive datasets LLPS and hsLLPS into training and test sets using various random selections, varying the training test size between 65-85%, and applied stratified sampling for the negative nsLLPS and hsnLLPS sets (**Table S2**). For parametrisation we removed the ‘uncharacterised’, ‘putative’ proteins and ‘coil-coiled’ domains from the nsLLPS and hsnLLPS datasets. Owing to their repetitive sequences coiled-coil domains still present a challenge for disorder predictions. We used the R program to determine the coefficients for the independent variables on the LLPS_train_ and hsLLPS_train_ datasets (**Table S5**). To decide the final coefficients, we aimed at high sensitivity as we expected many false positives in the negative datasets (proteins, which have not yet been reported to form droplets) and we aimed to find coefficients, which were consistent for many datasets.

The performance of the different models was evaluated based on area-under-the-curve (AUC), specificity, sensitivity and accuracy, which were computed by the R program (**Table S5**). The threshold for probability for droplet formation (*p*_*LLPS*_ => 0.61) was determined based on the logistic model, and were in good agreement for training and test sets. The π-π term was evaluated by the scripts given in the reference (19) using the same training and test sets (**Table S5**).

### Predicting the droplet state in different proteomes

The UniProt Swiss-Prot (reviewed) sequences were downloaded for *Caenorhabditis elegans, Drosophila melanogaster, Homo sapiens, Mus musculus, Rattus norvegicus, Saccharomyces cerevisiae, Schizosaccharomyces pombe* organisms and TrEMBL for *Xenopus laevis.* The degree of disorder was computed by the Espritz NMR program (36), the binding mode (p_DD_) was predicted for each residue using the FuzPred program (23). The probability of droplet formation for each protein was determined based on Eq. 3, with the coefficients given in **Table S5**. In each organism, we determined the frequency of proteins (including putative proteins), with *p*_*LLPS*_=> 0.6 (**Table S8**).

#### Observation of α-synuclein and β-synuclein liquid-liquid phase separation

Wild type α-synuclein and β-synuclein were purified from *E. coli* expressing plasmid pT7-7 encoding for the protein as previously described (38, 39). Following purification, the protein was concentrated using Amicon Ultra-15 Centrifugal Filter Units (Merck Millipore) and buffer exchanged into PBS at pH 8.0. Protein was subsequently labelled with 10-fold molar excess of Fluorescein 5-isothiocyanate (Sigma) for 3 hours at room temperature, followed by an overnight incubation at 4 °C with constant mixing. The excess dye was removed on a Sephadex G-25 desalting column (Sigma) and used immediately for phase separation experiments.

To induce droplet formation, non-labelled wild type α-synuclein and β-synuclein were mixed with FITC-labelled proteins at a 10:1 molar ratio in PBS with 50 mM NaCl and 10% polyethylene glycol (PEG) (Thermo Fisher Scientific). The final mixture was pipetted on a 35 mm glass-bottom dish (P35G-1.5-20-C, MatTek Life Sciences) and immediately imaged on a Leica TCS SP5 confocal microscope using a 40x/1.3 HC PL Apo CS oil objective (Leica Microsystems) with the temperature-controlled set at either 20 or 30 °C. The excitation wavelength was 488 nM for all experiments. All images were processed and analysed in ImageJ (NIH).

#### Complexin-1 phase separation

Recombinant human complexin-1 was obtained from NKmaxbio (Cat No: CPX0901). The C-terminal cysteine (C118) was labelled with 1.5X molar excess Alexa Fluor 488 C_5_ maleimide (Cat No: A10254, Life Technologies) overnight at 4 °C. The excess dye was removed on a Sephadex G-25 desalting column (Cat No: G25150-100G, Sigma) and the protein was buffer exchanged into 50 mM Tris-HCl (pH 7.4).

For imaging, 10 μM non-labelled complexin-1 was mixed with 10% (1 μM) Alexa Fluor 488 labelled protein in 50 mM Tris-HCl (pH 7.4), 100 mM NaCl, 1 mM DTT and 5% polyethylene glycol (PEG) (Cat No: B219555, Thermo Fisher Scientific) at 20 °C. The final mixture was pipetted on a 35 mm glass-bottom dish (Part No: P35G-1.5-20-C, MatTek Life Sciences) and immediately imaged on a Leica TCS SP5 using a 40x/1.3 HC PL Apo CS oil objective (Leica Microsystems). The excitation wavelength was 488 nM for all experiments. All images were analysed with ImageJ (NIH).

#### Fluorescence recovery after photobleaching

Fluorescence recovery after photobleaching (FRAP) was performed on the setup described above, under the same experimental conditions. Bleaching was done using the 488 nM laser at 50% intensity, to obtain ±50-60% photobleaching. Images were captured at 600 ms intervals, following a 1.8 s pre-bleach sequence and a 1.2 s bleach. Intensity traces of the bleached area were background corrected and normalised. A non-linear function of the recovery curve was fitted to obtain a relative recovery rate (Prism 8, GraphPad).

## Acknowledgement

M.F. acknowledges the financial support of HAS-11015.

## Author contributions

M.H. performed the *in vitro* experiments, wrote the experimental results; A.H. performed the statistical analysis and participated in development of the bioinformatics models; M.F. developed the methodology and supervised the computational work; M.V. supervised the experimental work; M.F. and M.V. conceptualized the work, analysed the results and wrote the paper.

## Supplemental Information

**Table S1. Datasets of proteins (REV, UNI, HTS, PSP, LPS-D, LPS-C** and the combined **LLPS** database**) reported to undergo liquid-liquid phase separation and overlaps between the different datasets***(separate .xls)*. **Dataset of droplet-promoting regions (DPRs).**

Columns: UniProt ID (relevant isoforms listed), UniProt entry name, protein name, gene names, length, organism, subcellular location, function, involvement in disease.

The overlap between LPS-D and LPS-C is displayed in the ‘O’ column of the LPS-D dataset (1 if the entry is present in both LPS-D and LPS-C, and 0 if only present in LPS-D).

In the overlap table, the lower triangle displays the number of overlapping entries in the whole dataset, the upper triangle shows the overlap between human proteins only. The population of the datasets all/human is shown in parenthesis.

**Table S2. Datasets of proteins, which were not reported to undergo liquid-liquid phase separation and used as a negative dataset***(separate .xls)*. Combined dataset for different organisms (nsLLPS) and human proteins only (hnsLLPS).

Columns: UniProt ID, UniProt entry name, protein name, gene names, length, organism, subcellular location, function, involvement in disease.

**Table S3. Statistical significances of the differences in the amino acid composition between the different datasets: LLPS and nsLLPS; LPS-D and LPS-C; LPS-C, nsLLPS***(separate .xls).* p values were computed by Kolgomorov-Smirnov test of the R program.

**Table S4 Training and performance of the logistic models to predict droplet-promoting propensity profiles.** The different sheets represent training on different droplet region lengths using different training group sizes. The different rows within each sheet were computed using different randomly chosen training sets. The selected models are highlighted by blue.

**Table S5. Training and performance of the logistic model to predict droplet-promoting propensities of proteins and performance of the logistic model with** π-π **term included** (*separate .xls)*. The terms used in each model are displayed and training fractions are indicated. Stratified sampling was applied.

**Table S6. Droplet-promoting propensities of proteins related with Alzheimer’s disease** (*separate .xls)*

**Table S7. Droplet-promoting propensities of proteins in the Swiss-Prot human proteome** (*separate .xls)*. The number of droplet-promoting regions (=> 25 residues) is also displayed.

**Table S8. Droplet-promoting propensities of proteins in***Caenorhabditis elegans, Saccharomyces cerevisiae, Schizosaccharomyces pombe, Drosophila melanogaster, Xenopus laevis, Mus musculus, Rattus norvegicus* (*separate .xls*). The frequency of droplet-forming proteins in different proteomes is displayed based on p_LLPS_ values and droplet-promoting regions with different lengths.

**Figure S1.**
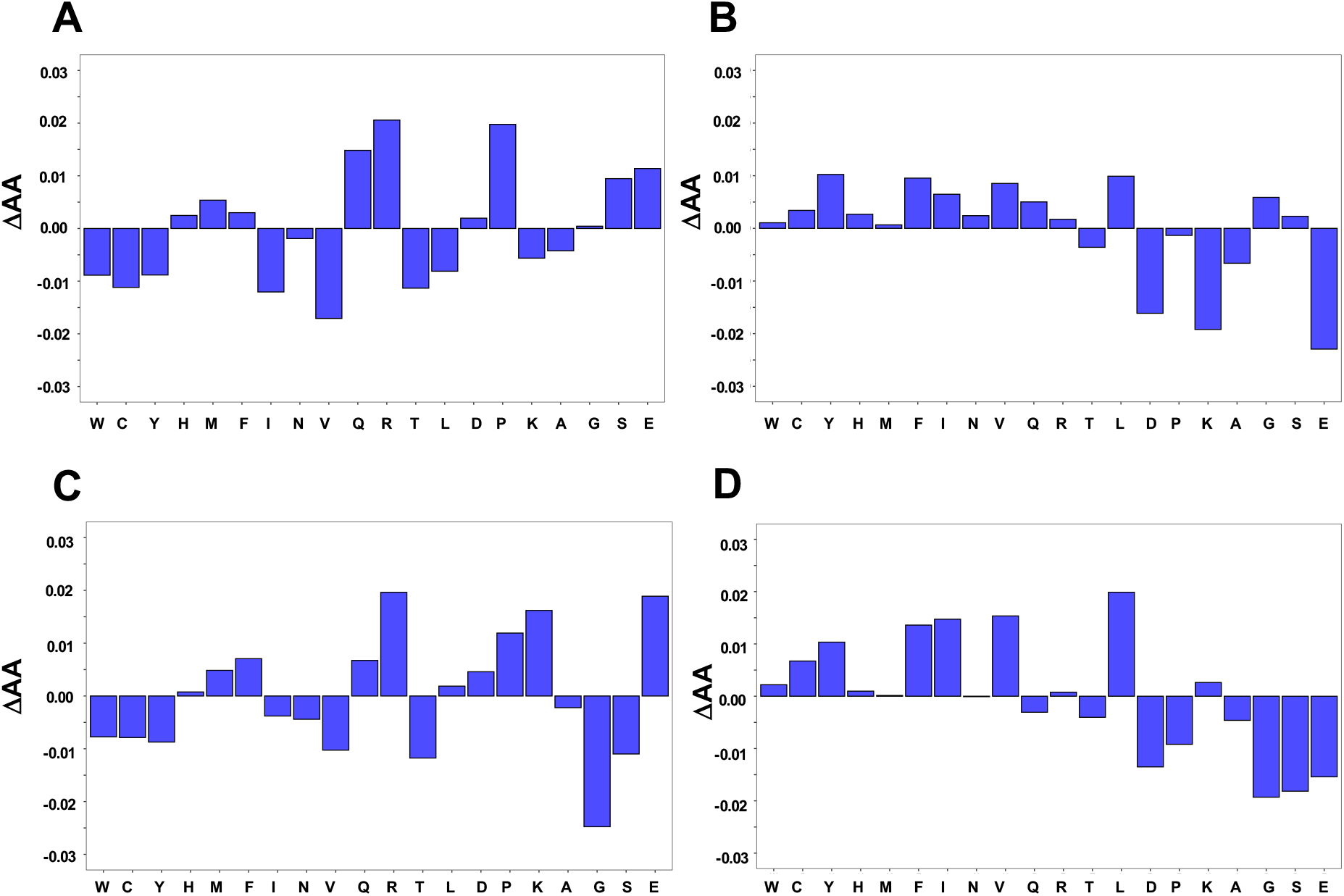
Differences in amino acid compositions of droplet-driving (A,B) and droplet-client (C,D) proteins from globular (A,C) and disordered proteins (B,D). Reference composition for disordered proteins was taken from the DisProt 7.0 database (34), that of globular proteins was taken from the reference (33). Amino acids are listed in increasing order in DisProt 7.0 database.

**Figure S2.**
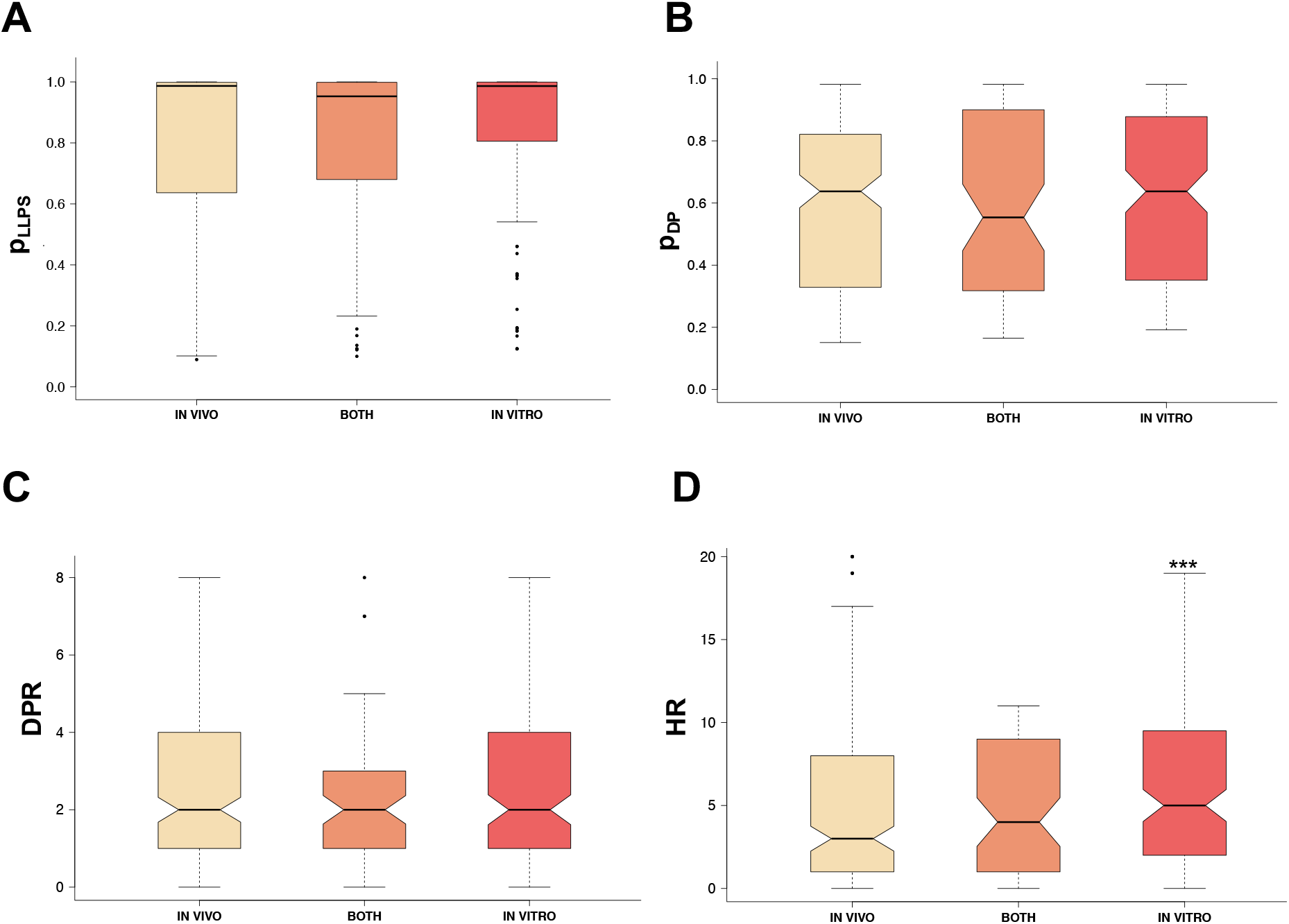
Droplet-promoting propensities of proteins forming droplets under physiological (in vivo, wheat), and non-physiological (in vitro, red) or both (orange) conditions. Experimental conditions were derived from the PhaSepDB dataset (http://db.phasep.pro) (29), using only reviewed (REV) evidences. **(A)** Droplet-promoting propensities of proteins (p_LLPS_), which form droplets under physiological and non-physiological conditions do not differ significantly. **(B)** The droplet-promoting propensity profiles (median p_DP_) and **(C)** the number of droplet-promoting regions (DPR) are also similar in proteins forming droplets under physiological and non-physiological conditions. **(D)** The number of hydrophobic motifs (HR) in disordered regions, however, is significantly enriched in proteins, which were observed to form droplets in vitro. Droplet-promoting propensities were computed by the FuzDrop program, statistical significances were determined by Mann-Whitney tests (***p < 10^−3^) using the R program.

**Figure S3.**
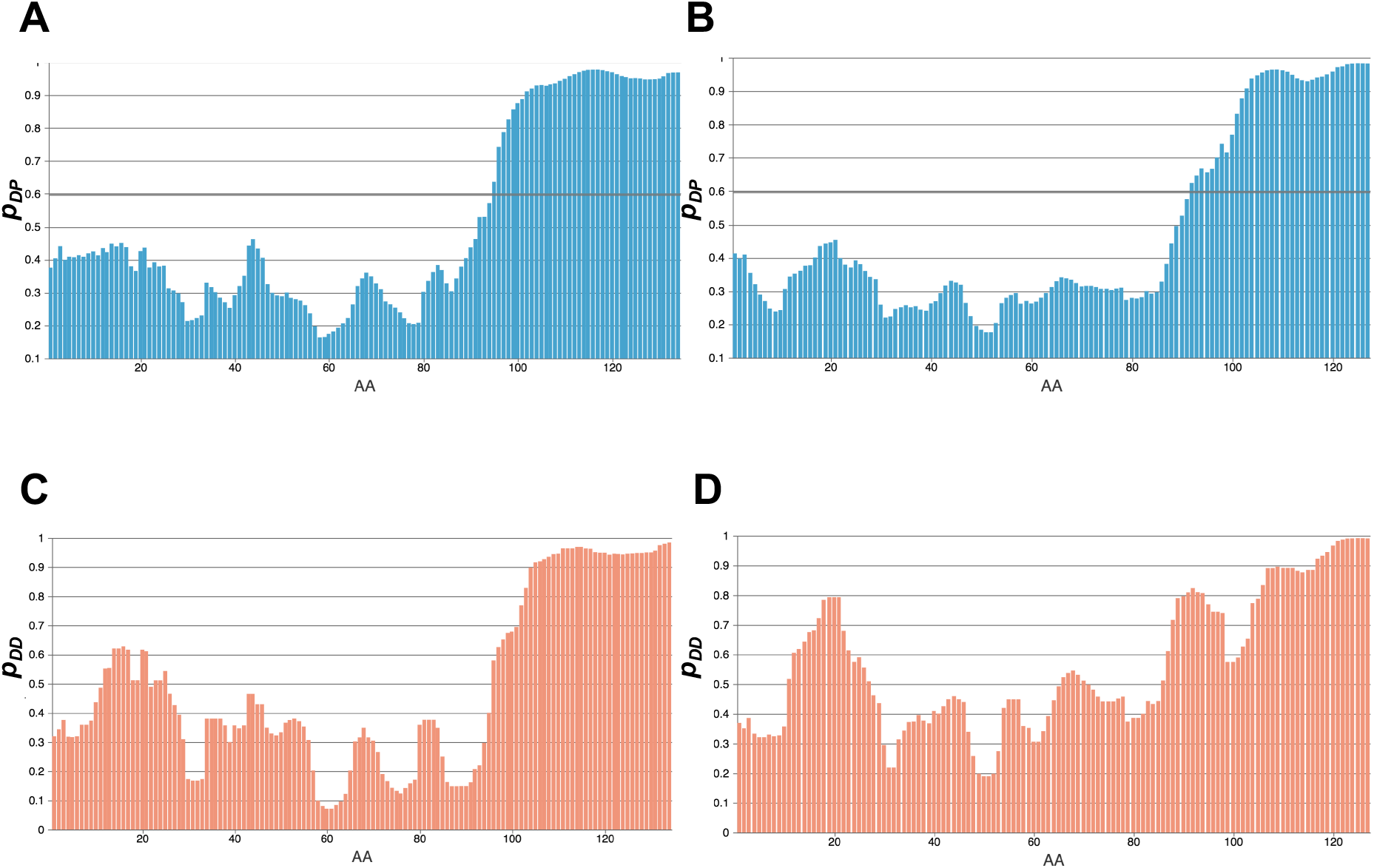
Droplet-promoting propensities (*p*_*DP*_) and binding entropies (*p*_*DD*_) of β-synuclein (A,C) and γ-synuclein (B,D). Both proteins, as well as α-synuclein, possess a droplet-promoting region in the C-terminus. The lack of a hydrophobic stretch in the NAC region, in contrast to α-synuclein, reduces the propensity for droplet formation. The predicted droplet forming propensities are 0.54 for β-synuclein and 0.40 for γ-synuclein. Droplet-promoting propensity profiles were computed by the FuzDrop, p_DD_ values by the FuzPred program (23).

**Figure S4.**
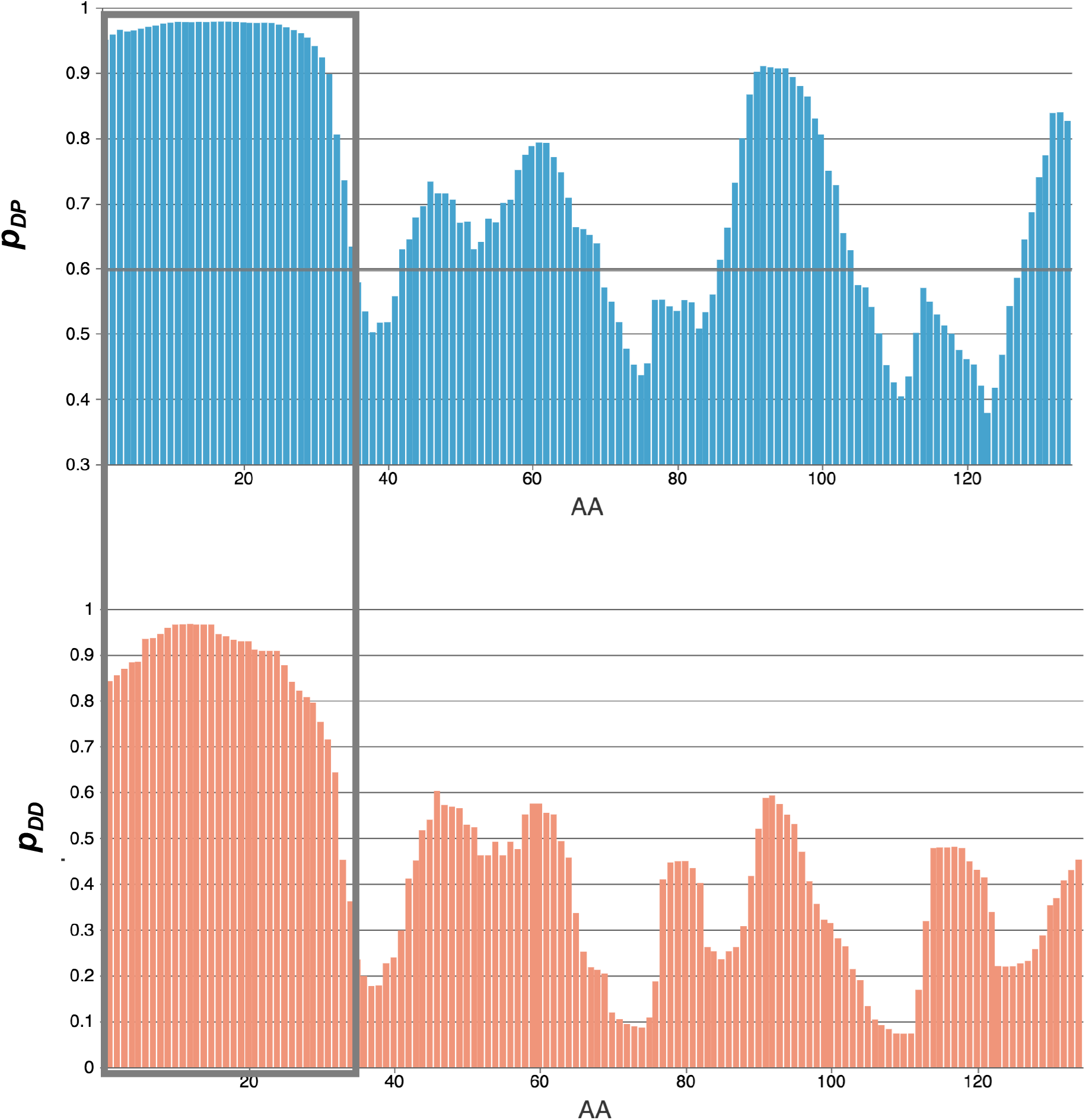
Prediction of droplet-promoting propensities (*p*_*DP*_) and probabilities for disordered binding (*p*_*DD*_) for complexin-1. The N-terminal region (residues 1-28, dark box) is predicted to be highly disordered in the bound state, in line with its high droplet-promoting propensity. In addition, most of the region, which interacts with the SNARE complex, also likely forms droplets.

**Figure S5.**
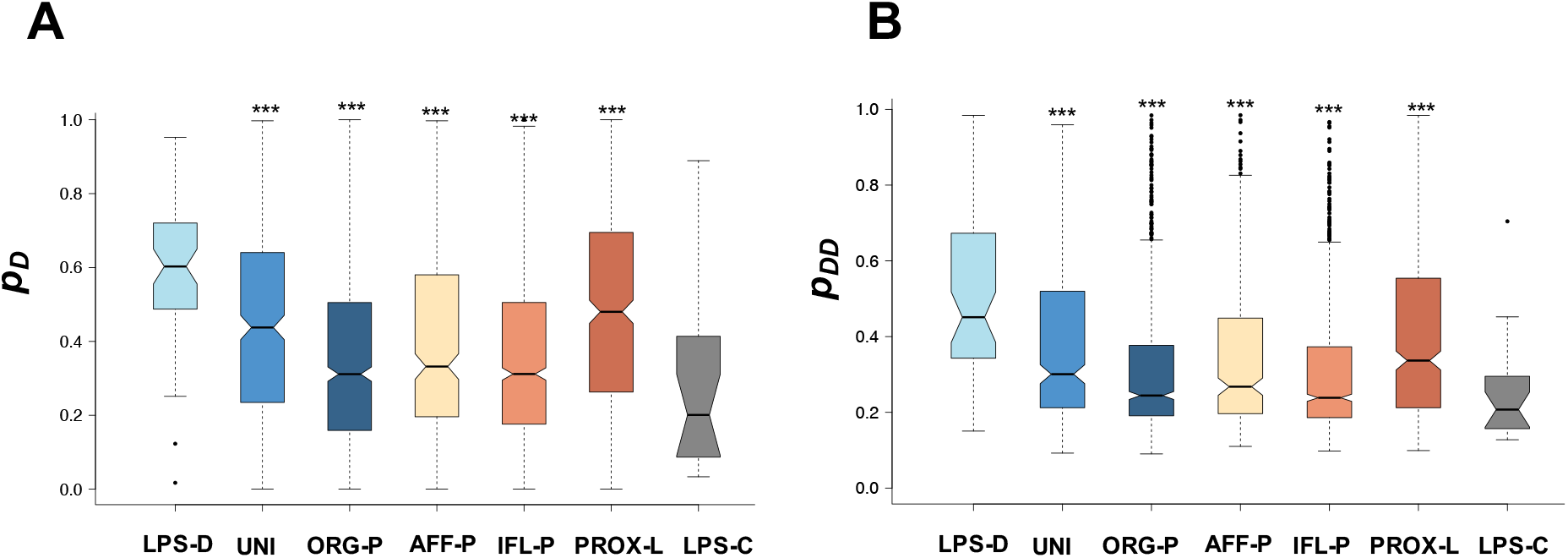
Comparison of probabilities for disorder in the free form (A) and disordered binding (B) in proteins in known human membraneless organelles (UNI) and proteins identified by high-throughput proteomic studies (Table S1) with droplet-driving (LPS-D) and droplet-client proteins (LPS-C). Proteins are grouped by the experimental technique: organelle purification (45, 46) (ORG-P, dark blue);affinity purification (47, 48) (AFF-P, wheat); immunofluorescence image based screen (49, 50) (IFL-L, orange); proximity labelling (51, 52) (PROX-L, brown). Proteins identified by proximity labeling have the largest fraction of components, which can spontaneously phase-separate. The probability for disorder estimates the conformational entropy in the free state (*p*_*D*_) based on the fraction of disordered residues using the Espritz NMR program (36). Probability for disordered binding estimates binding entropy based on the probability for disorder-to-disorder transition (p_DD_) using the FuzPred program (23). Based on these two characteristics, we estimate that human membraneless organelles are composed both by proteins that can spontaneously form droplets and by proteins that require additional partners for droplet formation. The statistical significances of the differences from LPS-D and LPS-C datasets were computed by Kolgomorov-Smirnov and Mann-Whitney tests (***p < 10^−3^) using the R program.

## Notes

### Competing Interest Statement

The authors have declared no competing interest.

## References

1. Hyman AA, Weber CA, & Julicher F (2014) Liquid-liquid phase separation in biology. Annu Rev Cell Dev Biol 30:39–58.

2. Banani SF, Lee HO, Hyman AA, & Rosen MK (2017) Biomolecular condensates: organizers of cellular biochemistry. Nat Rev Mol Cell Biol 18(5):285–298.

3. Bergeron-Sandoval LP, Safaee N, & Michnick SW (2016) Mechanisms and Consequences of Macromolecular Phase Separation. Cell 165(5):1067–1079.

4. Boeynaems S, et al. (2018) Protein Phase Separation: A New Phase in Cell Biology. Trends Cell Biol 28(6):420–435.

5. Yuan C, et al. (2019) Nucleation and Growth of Amino Acid and Peptide Supramolecular Polymers through Liquid-Liquid Phase Separation. Angew Chem Int Ed Engl 58(50):18116–18123.

6. Brangwynne CP, Tompa P, & Pappu RV (2015) Polymer physics of intracellular phase transitions. Nature Physics 11:899–904.

7. ten Wolde PR, Rein P, & Frenkel D (1999) Homogeneous nucleation and the Ostwald step rule. Physical Chemistry Chemical Physics 19.

8. Alberti S, Gladfelter A, & Mittag T (2019) Considerations and Challenges in Studying Liquid-Liquid Phase Separation and Biomolecular Condensates. Cell 176(3):419–434.

9. Sawaya MR, et al. (2007) Atomic structures of amyloid cross-beta spines reveal varied steric zippers. Nature 447(7143):453–457.

10. Knowles TP, et al. (2007) Role of intermolecular forces in defining material properties of protein nanofibrils. Science 318(5858):1900–1903.

11. Nott TJ, et al. (2015) Phase transition of a disordered nuage protein generates environmentally responsive membraneless organelles. Mol Cell 57(5):936–947.

12. Burke KA, Janke AM, Rhine CL, & Fawzi NL (2015) Residue-by-Residue View of In Vitro FUS Granules that Bind the C-Terminal Domain of RNA Polymerase II. Mol Cell 60(2):231–241.

13. Wang J, et al. (2018) A Molecular Grammar Governing the Driving Forces for Phase Separation of Prion-like RNA Binding Proteins. Cell 174(3):688–699 e616.

14. Hughes MP, et al. (2018) Atomic structures of low-complexity protein segments reveal kinked beta sheets that assemble networks. Science 359(6376):698–701.

15. Dignon GL, Best RB, & Mittal J (2020) Biomolecular Phase Separation: From Molecular Driving Forces to Macroscopic Properties. Annu Rev Phys Chem 71:53–75.

16. Krainer G, et al. (2020) Reentrant liquid condensate phase of proteins is stabilized by hydrophobic and non-ionic interactions. bioRxiv 2020.05.04.076299.

17. Vernon RM & Forman-Kay JD (2019) First-generation predictors of biological protein phase separation. Curr Opin Struct Biol 58:88–96.

18. Bolognesi B, et al. (2016) A Concentration-Dependent Liquid Phase Separation Can Cause Toxicity upon Increased Protein Expression. Cell reports 16(1):222–231.

19. Vernon RM, et al. (2018) Pi-Pi contacts are an overlooked protein feature relevant to phase separation. eLife 7.

20. Li P, et al. (2012) Phase transitions in the assembly of multivalent signalling proteins. Nature 483(7389):336–340.

21. Wu H & Fuxreiter M (2016) The Structure and Dynamics of Higher-Order Assemblies: Amyloids, Signalosomes, and Granules. Cell 165(5):1055–1066.

22. Hahn S (2018) Phase Separation, Protein Disorder, and Enhancer Function. Cell 175(7):1723–1725.

23. Miskei M, Horvath A, Vendruscolo M, & Fuxreiter M (2020) Sequence-Based Prediction of Fuzzy Protein Interactions. J Mol Biol 432:2289–2303.

24. Sormanni P, et al. (2017) Simultaneous quantification of protein order and disorder. Nat Chem Biol 13(4):339–342.

25. Fuxreiter M (2018) Fold or not to fold upon binding - does it really matter? Current Opinion in Structural Biology 54:19–25.

26. Kussie PH, et al. (1996) Structure of the MDM2 oncoprotein bound to the p53 tumor suppressor transactivation domain. Science 274(5289):948–953.

27. Horvath A, Miskei M, Ambrus V, Vendruscolo M, & Fuxreiter M (2020) Sequence-based prediction of protein binding mode landscapes PLoS Comp Biol 16:e1007864.

28. Leuenberger P, et al. (2017) Cell-wide analysis of protein thermal unfolding reveals determinants of thermostability. Science 355(6327).

29. You K, et al. (2020) PhaSepDB: a database of liquid-liquid phase separation related proteins. Nucleic Acids Res 48(D1):D354–D359.

30. Meszaros B, et al. (2020) PhaSePro: the database of proteins driving liquid-liquid phase separation. Nucleic Acids Res 48(D1):D360–D367.

31. Li Q, et al. (2020) LLPSDB: a database of proteins undergoing liquid-liquid phase separation in vitro. Nucleic Acids Res 48(D1):D320–D327.

32. Alberti S, Halfmann R, King O, Kapila A, & Lindquist S (2009) A systematic survey identifies prions and illuminates sequence features of prionogenic proteins. Cell 137(1):146–158.

33. Tompa P (2002) Intrinsically unstructured proteins. Trends Biochem Sci 27(10):527–533.

34. Hatos A, et al. (2020) DisProt: intrinsic protein disorder annotation in 2020. Nucleic Acids Res 48(D1):D269–D276.

35. Milles S, et al. (2015) Plasticity of an Ultrafast Interaction between Nucleoporins and Nuclear Transport Receptors. Cell 163(3):734–745.

36. Walsh I, Martin AJ, Di Domenico T, & Tosatto SC (2012) ESpritz: accurate and fast prediction of protein disorder. Bioinformatics 28(4):503–509.

37. Conicella AE, Zerze GH, Mittal J, & Fawzi NL (2016) ALS Mutations Disrupt Phase Separation Mediated by alpha-Helical Structure in the TDP-43 Low-Complexity C-Terminal Domain. Structure 24(9):1537–1549.

38. Ray S, et al. (2020) alpha-Synuclein aggregation nucleates through liquid-liquid phase separation. Nat Chem.

39. Hardenberg M, et al. (2020) Observation of an α-synuclein liquid droplet state and its maturation into Lewy body-like assemblies. BioRxiv.

40. Guenther EL, et al. (2018) Atomic structures of TDP-43 LCD segments and insights into reversible or pathogenic aggregation. Nat Struct Mol Biol 25(6):463–471.

41. Fusco G, et al. (2016) Structural basis of synaptic vesicle assembly promoted by alpha-synuclein. Nature communications 7:12563.

42. Lambert JC, et al. (2013) Meta-analysis of 74,046 individuals identifies 11 new susceptibility loci for Alzheimer’s disease. Nat Genet 45(12):1452–1458.

43. Lai Y, et al. (2016) N-terminal domain of complexin independently activates calcium-triggered fusion. Proc Natl Acad Sci U S A 113(32):E4698–4707.

44. Xue M, et al. (2010) Binding of the complexin N terminus to the SNARE complex potentiates synaptic-vesicle fusogenicity. Nat Struct Mol Biol 17(5):568–575.

45. Hubstenberger A, et al. (2017) P-Body Purification Reveals the Condensation of Repressed mRNA Regulons. Mol Cell 68(1):144–157 e145.

46. Andersen JS, et al. (2005) Nucleolar proteome dynamics. Nature 433(7021):77–83.

47. Ayache J, et al. (2015) P-body assembly requires DDX6 repression complexes rather than decay or Ataxin2/2L complexes. Mol Biol Cell 26(14):2579–2595.

48. Jonson L, et al. (2007) Molecular composition of IMP1 ribonucleoprotein granules. Mol Cell Proteomics 6(5):798–811.

49. Fong KW, et al. (2013) Whole-genome screening identifies proteins localized to distinct nuclear bodies. J Cell Biol 203(1):149–164.

50. Berchtold D, Battich N, & Pelkmans L (2018) A Systems-Level Study Reveals Regulators of Membrane-less Organelles in Human Cells. Mol Cell 72(6):1035–1049 e1035.

51. Markmiller S, et al. (2018) Context-Dependent and Disease-Specific Diversity in Protein Interactions within Stress Granules. Cell 172(3):590–604 e513.

52. Youn JY, et al. (2018) High-Density Proximity Mapping Reveals the Subcellular Organization of mRNA-Associated Granules and Bodies. Mol Cell 69(3):517–532 e511.

53. Shi M, Zhang P, Vora SM, & Wu H (2020) Higher-order assemblies in innate immune and inflammatory signaling: A general principle in cell biology. Curr Opin Cell Biol 63:194–203.

54. Fuxreiter M (2018) Fuzziness in Protein Interactions-A Historical Perspective. J Mol Biol 430(16):2278–2287.

55. Heller GT, Sormanni P, & Vendruscolo M (2015) Targeting disordered proteins with small molecules using entropy. Trends Biochem Sci 40(9):491–496.

56. Cremades N, et al. (2012) Direct observation of the interconversion of normal and toxic forms of α-synuclein. Cell 149(5):1048–1059.

57. Hoyer W, et al. (2002) Dependence of α-synuclein aggregate morphology on solution conditions. J. Mol. Biol. 322(2):383–393.

58. Vecchi G, et al. (2020) Proteome-wide observation of the phenomenon of life on the edge of solubility. Proc Natl Acad Sci U S A 117(2):1015–1020.

